# Cities of the future, visualizing climate change to inspire actions

**DOI:** 10.1101/458018

**Authors:** Jean-Francois Bastin, Emily Clark, Thomas Elliott, Simon Hart, Johan van den Hoogen, Iris Hordijk, Haozhi Ma, Sabiha Majumder, Gabriele Manoli, Julia Maschler, Lidong Mo, Devin Routh, Kailiang Yu, Constantin Zohner, Thomas, W. Crowther

## Abstract

Combating against climate change requires unified action across all sectors of society. However, this collective action is precluded by the ‘consensus gap’ between scientific knowledge and public opinion. A growing body of evidence suggests that facts do not persuade people to act. Instead, it is visualization - the ability to simulate relatable scenarios - is the most effective approach for motivating behavior change. Here, we exemplify this approach, using current climate projections to enable people to visualize cities of the future, rather than describing intangible climate variables. Analyzing city pairs for 520 major cities of the world, we characterize which cities will most closely resemble the climate conditions of which other major cities by 2050. On average, most cities will resemble cities that are over 1000km south, and 22% of cities will experience climate conditions that are not currently experienced by any existing major cities. We predict that London’s climate in 2050 will resemble Barcelona’s climate today, Madrid will resemble to Marrakesh, Moscow to Sofia, Seattle to San Francisco, Stockholm to Budapest, Tokyo to Changsha, etc. Our approach illustrates how complex climate data can be packaged to provide tangible information. By allowing people to visualize their own climate futures, we hope to empower citizens, policy makers and scientists to visualize expected climate impacts and adapt decision making accordingly.

---

Climate change is a systemic threat to humankind ^1^. In 1992, 172 nations convened in Rio de Janeiro to establish the United Nations Framework Convention on Climate Change (UNFCCC), recognizing the urgent need for climate action. Yet, 26 years later, we still have to alter the course of this global threat ^2^. The UN estimates that the cost of climate change under a business as usual scenario will exceed $12 trillion by 2050 ^3^. Less than 0.1% of this amount has been committed to the UNFCCC’s Green Climate Fund to incentivize global climate action ^4^. Despite the overwhelming scientific consensus that human activity is driving climate change ^1,5^, 53% of people in the world still do not believe that climate change will affect them during their lifetime ^6^.

The gap between scientific and public understanding (referred to as the Consensus Gap) is largely attributed to failures in climate change communication ^7^. Often limited to ad-hoc reporting of extreme weather events or intangible, long-term climate impacts (e.g. changes in average temperature by 2100). The intangible nature of this reporting fails to adequately convey the urgency of this issue to a public audience on a consistent basis ^8^. It is hard for most people to envision how an additional 2°C of warming might affect daily life. This ineffective communication of climate change facts, compounded by the uncertainty about the extent of expected changes, has left the door open for widespread misinterpretation about the existence of this unassailable global phenomenon.

History has repeatedly shown us that data and facts alone do not inspire humans to change their beliefs or act ^8^. Increased scientific literacy has no correlation with the acceptance of climate change facts ^9^, and can instead have an opposite effect. A growing body of research demonstrates that visualization - the ability to create a mental image of the problem - is the most effective approach for motivating behavior change ^10,11^. The greatest challenge facing climate scientists is no longer the detailed characterization of future climate, but the development of communication mechanisms that inspire coordinated action. We argue that enabling people to visualize a tangible climate future is the most crucial objective facing current climate science.

Here, we implement this strategy, taking the current climate data for the world’s major cities (Current Cities), and projecting what they will be like in 2050 (Future Cities). Rather than describing the quantitative changes in climate variables, we quantify which Current Cities will most closely resemble the climate conditions of Future Cities (see Material and Methods). For example, using our approach we can predict that London’s climate in 2050 will resemble Barcelona’s climate today. Our approach makes the intangible (e.g. a given rise in temperature) tangible, in terms of people’s current experience. This is intended to capture people’s imagination, not only because over 68% of the world’s population will live urban areas ^12^, but also because iconic locations allow us to visualize the imminent changes that are likely to occur within our lifetimes.

We characterized the climate of the world’s 520 major cities using 19 climatic variables that reflect the variability in temperature and precipitation regimes for current and future conditions. Future conditions are estimated using a realistic Representative Concentration Pathway (RCP4.5), which considers a stabilization of CO2 emissions by mid-century (see Material and Methods). Using a multivariate analysis, we analyzed the climate similarity of all Current and Future cities to one another (Figure 1, Database S1). This simple analysis enable us to estimate which major cities of the world will remain relatively similar, and which will shift to reflect the climate of another city by 2050. Overall, our analysis shows that 77% of the world’s Current Cities will experience a striking change in climate conditions, making them more similar to the conditions of another existing city (see Databases S1-S2). In addition, 22% of cities will experience climate conditions that are unlike any major city currently existing on the planet (Figure 2a-b).

**Figure 1.**
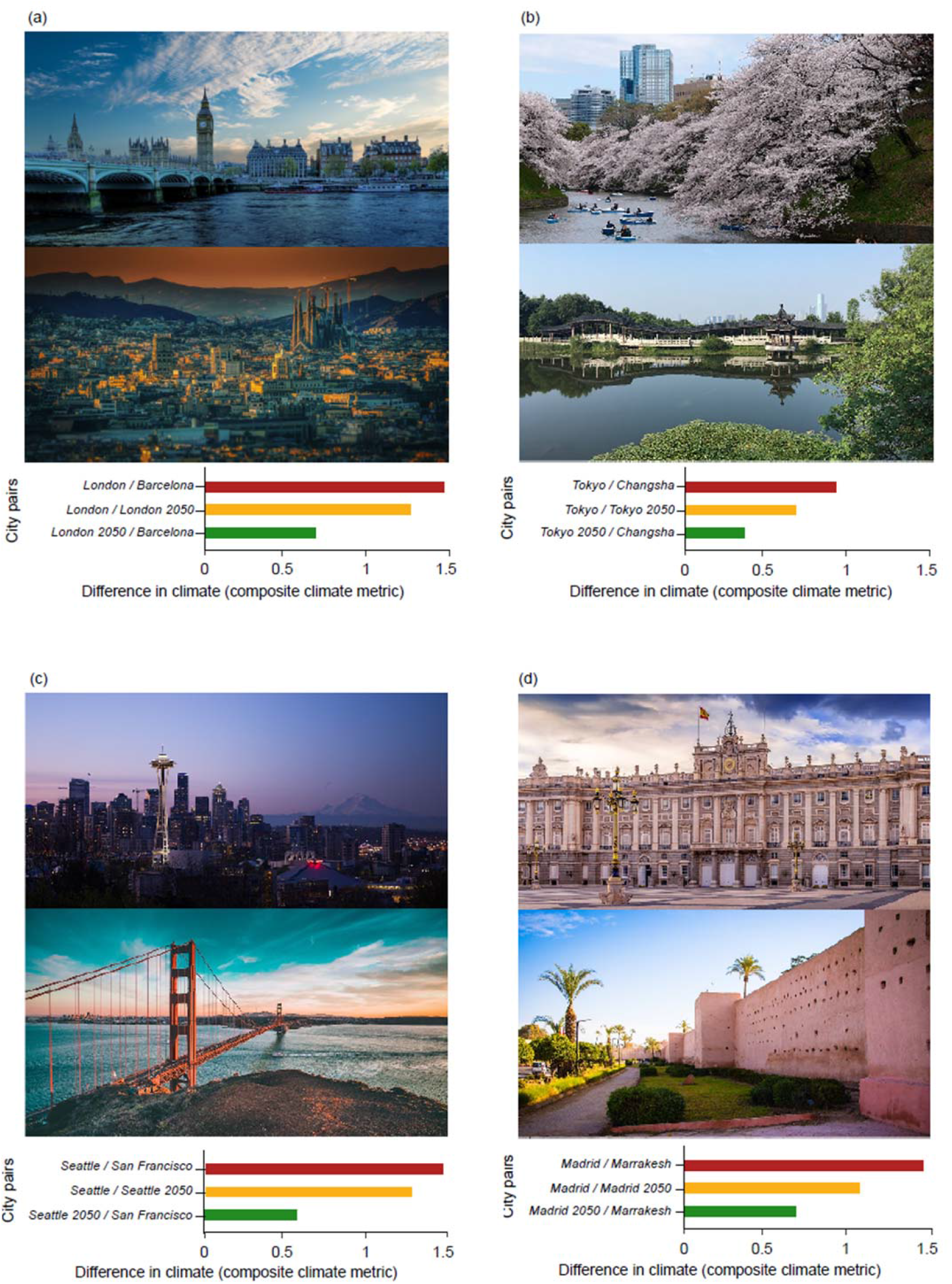
Future cities and similar current climate counterpart. Difference between future and current climate for four cities and an example of their similar current counterpart. Illustration of the results of the analysis for London (**a**; counterpart: Barcelona), Buenos Aires (**b**; counterpart: Sidney), Nairobi (**c**; counterpart:Beirut) and Portland (**d**; counterpart:San Antonio). Images of Barcelona and London were obtained on Pixabay, shared under common creative CC0 license.

**Figure 2.**
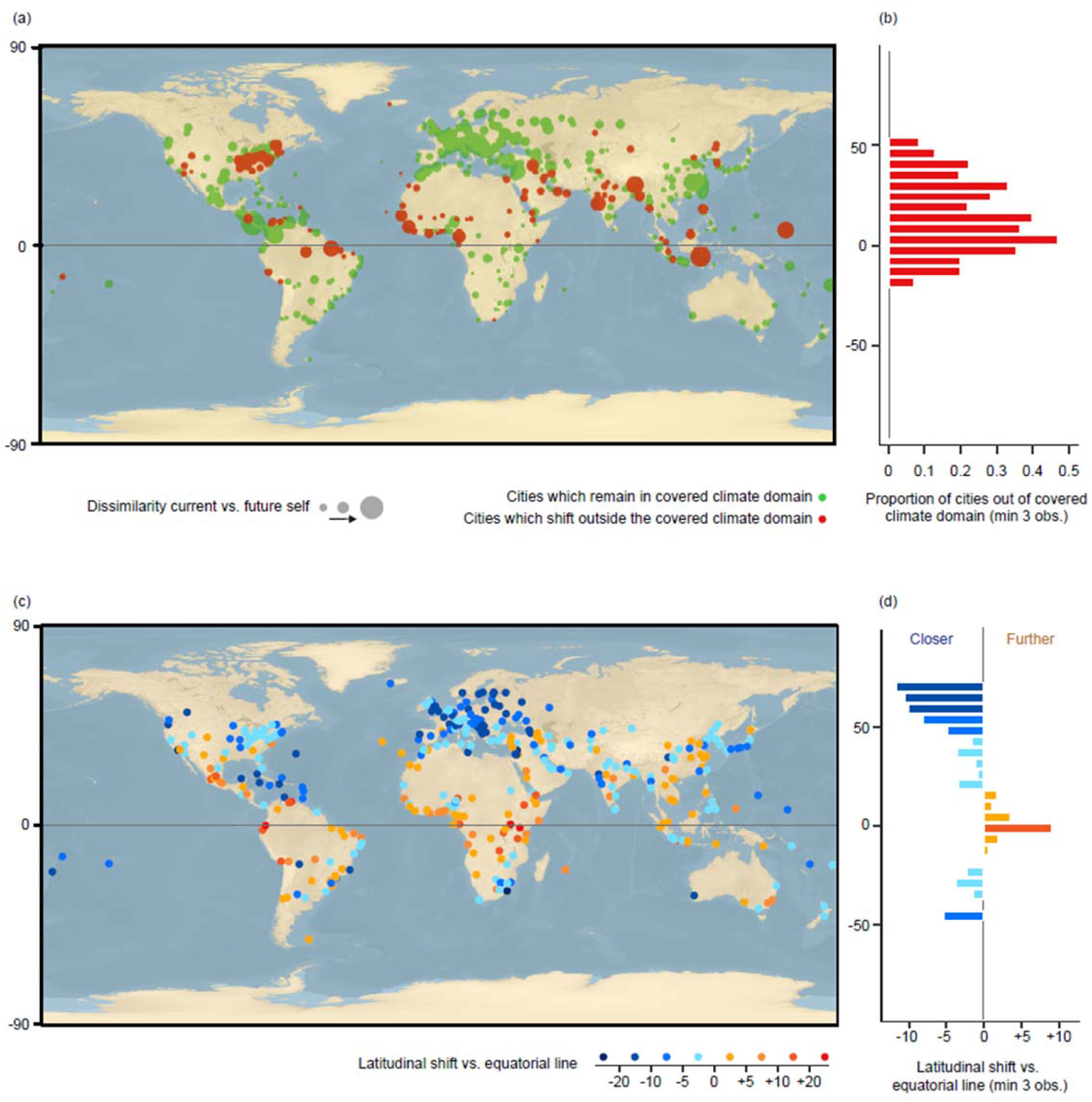
Extent of climate changes in major cities of the world by 2050. **a, b**, the extent of change in climate conditions. Cities predicted to have climates that no major city has experienced before are colored in red (mostly within the tropics). Cities for which future climate conditions reflect current conditions in other major cities of the world are shown in green. The size of the dots represents the magnitude of change between current and future climate conditions. **b**, The proportion of cities shifting away from the covered climate domain (concentrated in the tropics). **c,d**, The extent of latitudinal shifts in relation to the equatorial line. Cities shifting towards the equator are colored with a blue gradient (mostly outside the tropics), while cities shifting away from the equator are colored with a yellow to red gradient (mostly within the tropics). **d**, A summary of the shift by latitude is illustrated in a barchart, with shifts averaged by bins of 5 degrees

The proportion of shifting cities varied consistently across the world. Cities in northern latitudes will experience the most dramatic shifts in extreme temperature conditions (Figure 2c-d). For example, across Europe, both summers and winters will get warmer, with average increases of 3.5^°^ C and 4.7^°^ C, respectively. These changes would be equivalent to a city shifting 1,000 km further south, under current climate conditions (Figure 2c-d). Consequently, by 2050, striking changes will be observed across the northern hemisphere: London’s climate in 2050 will be more similar to the current climate in Barcelona than to London’s climate today; Stockholm will be more similar to Vienna; Moscow to Sofia; Portland to San Antonio, San Francisco to Lisbon and Tokyo to Changsha; (Figure 1, Database S2).

Cities in the tropical regions will experience smaller changes in average temperature, relative to the higher latitudes. However, shifts in rainfall regimes will dominate the tropical cities. This is characterized by both increases in extreme precipitation events (+5% rainfall wettest month) and, the severity and intensity of droughts (−14% rainfall driest month). With more severe droughts, tropical cities will move slightly away from the equator, towards drier climates (Figure 2c-d). However, the fate of major tropical cities remains highly uncertain because many tropical regions will experience unprecedented climate conditions. Specifically, of all 22% of cities that will experience novel climate conditions, most (64%) are located in the tropics. These include Manaus, Libreville, Kuala Lumpur, Jakarta, Rangoon, and Singapore (Figure 2a-b, Database S2).

Our analysis allows us to visualize a tangible climate future of the world’s major cities. These results enable decision makers from all sectors of society, to envision changes that are likely to occur in their own city, within their own lifetime. Londoners, for example, can start to consider how their 2050 equivalents (e.g. Barcelona today) have taken action to combat their own environmental challenges. In 2008, Barcelona experienced extreme drought conditions which required the importation of €22m of drinking water. Since then, the municipal government has implemented a series of ‘smart initiatives’ to manage the city’s water resources (including the control of park irrigation and water fountain levels). The Mayor of London has factored drought considerations into his Environment Strategy aims for 2050 ^13^, but this study can provide the context to facilitate the development of more targeted climate strategies. In addition, this information can also empower local citizens to evaluate proposed environmental policies. By allowing people to visualize their own climate futures, we hope that this information can engage people to take action, both to mitigate and adapt to climate change.

Our study is not a novel model revealing updated climate projections or expectations by 2050. Instead, our analysis is intended to illustrate how complex climate data can be effectively summarized into tangible information that can be easily interpreted by anyone. Of course, the climate scenarios that we have used are based on predictions from a few climate models, run under a single (business as usual) climate scenario. We recognize that these models are characterized by huge amounts of uncertainty ^14^, and the predicted Future Cities may change as these Earth System Models are refined, in particular in light of urban climate specificities ^15^. However, our results are likely to reflect the qualitative direction of climate changes within cities and so meet our primary goal, which is to communicate predicted climate changes to a non-specialist audience in order to motivate action. When model projections are updated, we would recommend communicating any new results with this goal in mind.

Finally, our work is a call to encourage scientists to take more responsibility in the fight against climate change. In addition to furthering current knowledge, we should improve our communication in order to empower stakeholders and policy makers that are trying to change the course of climate change. It is time to bridge the gap between knowledge and action.

## Materials and Methods

### Major cities of the world

We selected the “major” cities of the world from the “LandScan (2016) High Resolution global Population Data Set” created by the Oak Ridge National Laboratory^16^. By “major” cities, we considered cities that are an administrative capital or that account more than 1,000,000 inhabitants. In total, 520 cities were selected.

### The climate database

To characterize the current climate conditions among these major cities of the world, we extracted 19 bioclimatic variables from the latest Worldclim global raster layers (Version 2; period 1970-2000) at 30 arc-seconds resolution^17^. These variables captured various climatic conditions, including yearly averages, seasonality metrics, and monthly extremes for both precipitation and temperature at every location.

### Future data: GCMs, downscaling and future scenarios

For the future projections, the same 19 bioclimatic variables were averaged from the outputs of three general circulation models (GCM) commonly used in ecology^18,19^. Two Community Earth System Models (CESMs) were chosen as they investigate a diverse set of earth-system interactions: the CESM1 BGC (a coupled carbon–climate model accounting for carbon feedback from the land) and the CESM1 CAM5 (a community atmosphere model)^18^. Additionally, the Earth System component of the Met Office Hadley Centre HadGEM2 model family was used as the third and final model^19^. To generate the data, we chose Representative Common Pathway 4.5 (RCP 4.5) scenario from the Coupled Model Intercomparison Project Phase 5 (CMIP5) as the input. It is a stabilization scenario, meaning that it accounts for a stabilization of radiative forcing before 2100, anticipating the development of new technologies and strategies for reducing greenhouse gas emissions^20^. For each output, a delta downscaling method developed by the CGIAR Research Program on Climate Change, Agriculture and Food Security (CCAFS) was applied to reach a resolution of 30 arc-seconds^21^, using current conditions Worldclim 1.4 as a reference.

### Summarizing the current climate among the major cities through a principal component analysis

The 19 current and future bioclimatic variables were extracted from the coordinates of the 520 major cities (i.e., the city centroids), meaning each city had two sets of bioclimatic metrics: the current climate data for the world’s major cities (Current Cities) and the equivalent 2050 projection (Future Cities) according to the average of the three RCP 4.5 GCMs.

A principal components analysis (PCA) was then performed on current bioclimatic data in order to account for correlation between climate variables and to standardize their contributions to the subsequent dissimilarity analysis^22^. As the first four principal components accounted for more than 85% of the total variation of climate data (40.2%, 26.9%, 10.5% and 7.6%, respectively), the remaining principal components were dropped from later analyses. The main contributing variables to the four components are the temperature seasonality (axis 1), the minimum temperature of the coldest month (axis 1), the maximum temperature of the warmest month (axis 2), the precipitation seasonality (axis 2), the precipitation of the driest (axis 4) and of the wettest (axis 3) month, and the temperature diurnal range (axis 4, supplementary figure 1).

### Analysis of changes between current and future cities from the PCA

The future climate of each city was then projected within these four principal components (using the PCA eigenvectors derived from the bioclimatic variables of the current climate) to allow for direct comparison between Current and Future Cities (supplementary figure 1a-b). On the plane defined by the first two components of the PCA (supplementary figure 1a), we observe changes towards less temperature seasonality, with higher maximal and minimal temperatures during the year, as well as higher precipitation seasonality, with higher precipitation in the wettest month but lower precipitation in the driest one. While no clear trend can be observed along the third axis, the changes along the fourth axis show higher temperature diurnal range (supplementary figure 1b), i.e. the daily difference between cities’ maximum and minimum temperatures will increase. In brief, cities of the world become hotter, in particular during the winter and the summer. Wet seasons become wetter and dry season drier.

### Calculating the extent of the covered climate domain

For further interpretation of the results, a convex hull was computed from the coordinates of the Current Cities within the multivariate space defined by the first four principal components axes^23^. For reference, a convex hull of a set of N-dimensional points forms the smallest possible hypervolume (in N-dimensions) containing all points defined in that set; in this case, it defines the bounds of climatic combinations that Earth currently experiences in these 520 cities. All Future Cities falling outside the hypervolume of this convex hull represent currently non-existent bioclimatic assemblies in these cities. Overall 78% of the 520 Future Cities present a climate within this hypervolume, while 22% of the Future Cities’ climate conditions (many of which are distributed around the equator and in arid or semi-arid regions) would disappear from this current climatic domain (Figure 2).

### Pairing cities based on the similarity between current and future climate conditions

Euclidean distances (i.e., dissimilarity indices) were calculated for every combination of Current and Future City based on their coordinates within the multivariate space defined by the first four principal components axes, creating a symmetric dissimilarity matrix with pairwise comparisons for all cities (see Database S1). The Euclidean distance was calculated using the vegan package on R ^22^. Each Future City was then paired with its three closest Current Cities based on the dissimilarity values (see Databases S1-2). Three cities are kept for each Future city in order to facilitate comparison between Current and Future climate, as all cities are not necessarily known by the reader. To avoid shifts due to pixel mismatch between Current and Future climate conditions, the final analysis was performed keeping shift values between the 5^th^ and the 95^th^ percentile, i.e. keeping 477 out of the original 520 cities. Results show that 77% of the world’s Current Cities will experience a striking change in climate conditions, making them more similar to the conditions of another currently existing city.

### Calculating the standardized latitudinal shift

To illustrate and summarize the shifts between Current and Future Cities, we calculated the importance of latitudinal shift between the most similar pairs. Shifts in latitude were standardized for both hemisphere, so that a shift south in the northern hemisphere is equal to a shift north in the southern hemisphere. In other words, the standardized latitudinal shift illustrates a shift in relation to the equatorial line (shifting away from or towards the equator). The results (Figure 2) show that cities in boreal and temperate regions tend to move towards the equator while cities in tropical regions tend to move away from the equator.

Analyses and figures were performed using Rcran version 3.3.2, maps were built using Q-GIS 3.0.

## Acknowledgments

This work was supported by grants to T.W.C. from DOB Ecology, Plant-for-the-Planet and the German Federal Ministry for Economic Cooperation and Development. Images of cities were obtained on Pixabay, and openly shared under CC0 common creative license.

## Author Contributions

J.-F.B. conceptualized the study and did the analyses. J.-F.B., E.C., T.E., S.H., J.H, I.H., H.M., S.M., G.M., J.M., L.M., D.R., K.Y., C.Z. and T.W.C. wrote of the manuscript.

## Competing interests

The authors declare no competing interests.

## Material and Correspondance

Correspondence and material requests should be addressed to Jean-Francois Bastin

